# Survival intravascular photoacoustic imaging of lipid-rich plaque in cholesterol fed rabbits

**DOI:** 10.1101/2022.06.07.494321

**Authors:** Yi Zhang, Erik Taylor, Nasi Huang, James Hamilton, Ji-Xin Cheng

## Abstract

Intravascular photoacoustic (IVPA) imaging is a promising modality for quantitative assessment of lipid-laden atherosclerotic plaques. Yet, survival IVPA imaging of the same plaque in the same animal is not demonstrated. Here, using a sheathed IVUS/PA catheter of 0.9 mm in diameter, we demonstrate MRI-guided survival IVPA imaging of same plaque in an aorta of a well-established rabbit model mimicking atherosclerosis in human patients. The IVUS/PA results were confirmed by histology. These advances open the opportunity to evaluate the effectiveness of a therapy that aims to reduce the size of atherosclerotic plaques and demonstrates the potential of translating the IVPA catheter into clinic for detection of vulnerable plaques that are at high risk for thrombosis.

## Introduction

Atherosclerotic cardiovascular disease is the leading cause of mortality in the United States, causing nearly one million deaths every year^1,2^. Cardiovascular disease deaths are most often triggered by the rupture of vulnerable atherosclerosis plaques and subsequent thrombosis^3^. Early identification of vulnerable plaques in vivo, by characterization of morphological and compositional features of the plaque, can have a positive impact on guiding the treatment^4,5^. In human, atherosclerosis is initiated by the sub-endothelial infiltration of plasma cholesterol and cholesteryl ester, which is enhanced by endothelial damage and further promotes inflammation over decades, together leading to the plaque instability^6-8^. The size and location(s) of the lipid pool and the thickness of the fibrous cap are two features that can be used to determine the plaque stability^9,10^.

To diagnose the pathological stage of atherosclerosis reliably and accurately, both the morphological and chemical information of the plaque are required. Magnetic resonance imaging (MRI) is widely used to characterize major components of a plaque, including the fibrous cap, the lipid core hemorrhage and inflammation^11,12^. Nevertheless, non-invasive MRI is limited by the millimeter-scale spatial resolution. Several catheter-based invasive methods, including intravascular ultrasound (IVUS)^13,14^ and optical coherence tomography^15^, were developed to obtain high resolution morphological and location information about the plaques, yet lacking information about the chemical composition. IVUS was recently combined with near-infrared spectroscopy (NIRS), which provides information about the lipid content in a plaque^16^. However, NIRS cannot precisely locate the plaque and determine the volume of the lipid core.

Catheter-based intravascular photoacoustic (IVPA) imaging is a promising technology for localization, characterization, and quantification of lipids in a coronary atherosclerotic plaque. IVPA imaging is based on molecular absorption of a nanosecond laser pulse and subsequent generation of an ultrasound signal detected by a transducer; the method is capable of mapping the composition of the artery wall with an adequate imaging depth. Catheter-based IVPA imaging is minimally invasive, and provides high resolution chemical maps of the arterial wall ^17-19^. Co-registered with IVUS, the IVPA imaging system can achieve real time display of lipid-rich plaques with high sensitivity and sufficient depth, providing size, location, and composition information to identify plaques having high risk of rupture^19,20^.

Vibrational photoacoustic imaging of vascular lipids was first demonstrated by using a laser at wavelength around 1200 nm^21,22^ and then by a laser at 1700 nm^23,24^. Since then, there have been many advances in developing an IVPA catheter for intravital imaging of vascular lipids. Specifically, the Ji-Xin Cheng group demonstrated detection of lipid-laden plaque using a KTP-based OPO at 1.7 μm^25^, high-sensitivity IVPA imaging through a collinear catheter design^26^ and a portable IVUS-PA system capable of imaging at up to 25 frames per second in real-time display mode^27^. The Cheng group also demonstrated in vivo IVPA imaging of lipid distribution in rabbit aorta under clinically relevant conditions^28^, and more recently, a dual-transducer catheter for high-resolution IVUS and high-sensitivity IVPA imaging^29^. The Gijs van Soest group demonstrated IVPA imaging of human coronary atherosclerotic plaque ex vivo^30^, real-time volumetric lipid imaging in vivo at speed of 20 frames per second^31^, and lipid detection in a human atherosclerotic lesion at 1.7 μm with a lower pulse energy than that at 1.2 μm^32^. The Qifa Zhou and Zhongping Chen groups demonstrated high resolution IVPA imaging by using a transducer with a working frequency as high as 80 MHz^33^. The Stanislav Emelianov group demonstrated intravascular photoacoustic imaging of lipid in atherosclerotic plaques in the presence of luminal blood^24^. The same group also built a complete IVUS/IVPA imaging system capable of real-time IVUS/IVPA imaging, with online data acquisition, image processing, and display of both IVUS and IVPA images^34^. The Liang Song group demonstrated optical-resolution IVPA imaging offering optical-diffraction limited transverse resolution as fine as 19.6 μm^35^. The Song group also improved an IVPA/IVUS system to an imaging speed as high as 100 f/s based on a miniature catheter of 0.9 mm ^36^. The Da Xing Group demonstrated ex vivo and in vivo IVPA imaging of aortic plaques in rabbits^37^. The Shihe Yang group developed a tapered fiber-based IVPA catheter for high-resolution imaging of lipid-rich plaque^38^.

Despite these advances, to our knowledge, survival IVPA imaging on the same plaque in the same animal with atherosclerosis has not been conducted. Without such capability, the potential of using IVPA imaging to monitor plaque progression under different diets or drug treatment cannot be fully appreciated. Here, we demonstrate survival IVPA imaging of atherosclerotic plaques using a rabbit arterial wall balloon injury model. We successfully conducted in vivo IVPA imaging on same rabbits with survival surgeries. This advancement provides us with a new platform towards a deeper understanding of the atherosclerosis and an accurate way of evaluating the effectiveness of a treatment. Detailed methods and results are shown below.

## Models and Methods

### The rabbit model

New Zealand White (NZW) rabbits are used as an animal model to study atherosclerosis and sudden thrombosis in a controlled manner ^39^. These rabbits develop foam-cell-rich (fatty steaks) plaques when short term high-fat diets were the only stimulus used to induce atherosclerosis. Intermittent cycles of fat feeding with periods of normal diet induces plaques at more advanced stages that resembles human atheroma. Moreover, with the combination of arterial wall balloon injury and hyperlipidemia, advanced lesions form in shorter periods (∼6 weeks) compared to the cholesterol-fed swine models (∼ 4 months)^40^.

Using this rabbit model, a recent MRI study of atherosclerosis monitored plaque progression preceding stimulated plaque rupture with serial analysis in the same animal^41^. Each aorta typically has several plaques, each of which can be monitored over time, and the stable and vulnerable plaques can be differentiated at the end of the 3-month protocol by triggering. Notably, the size of its aorta is similar to a human coronary artery^39,42^. These unique features render this rabbit model an ideal testbed for survival in vivo IVPA imaging. In our experiment, two 1-month old male NZW rabbits were chosen as the test group and one 1-month male NZW rabbit was chosen as the control group. The experimental design is shown in **Figure 1**. The test group was on 1% cholesterol diet for 2 weeks before balloon surgery, followed by 1% cholesterol diet for another 6 weeks, then on normal chow diet for 4 weeks. The control group was on normal chow diet all 12 weeks. The imaging procedure was the same for both groups. In vivo MRI was first conducted to locate the plaque. Four weeks after MRI, survival IVPA imaging was conducted to map the lipids inside the plaque. Then, rabbits were allowed to recover during 4 to 6 weeks. Next, plaque disruption was attempted by two pharmacological triggers on two consecutive days. After the triggering, second IVPA imaging was conducted. Finally, the rabbits were euthanized and aorta tissues were extracted for histology examination.

**Figure 1.**
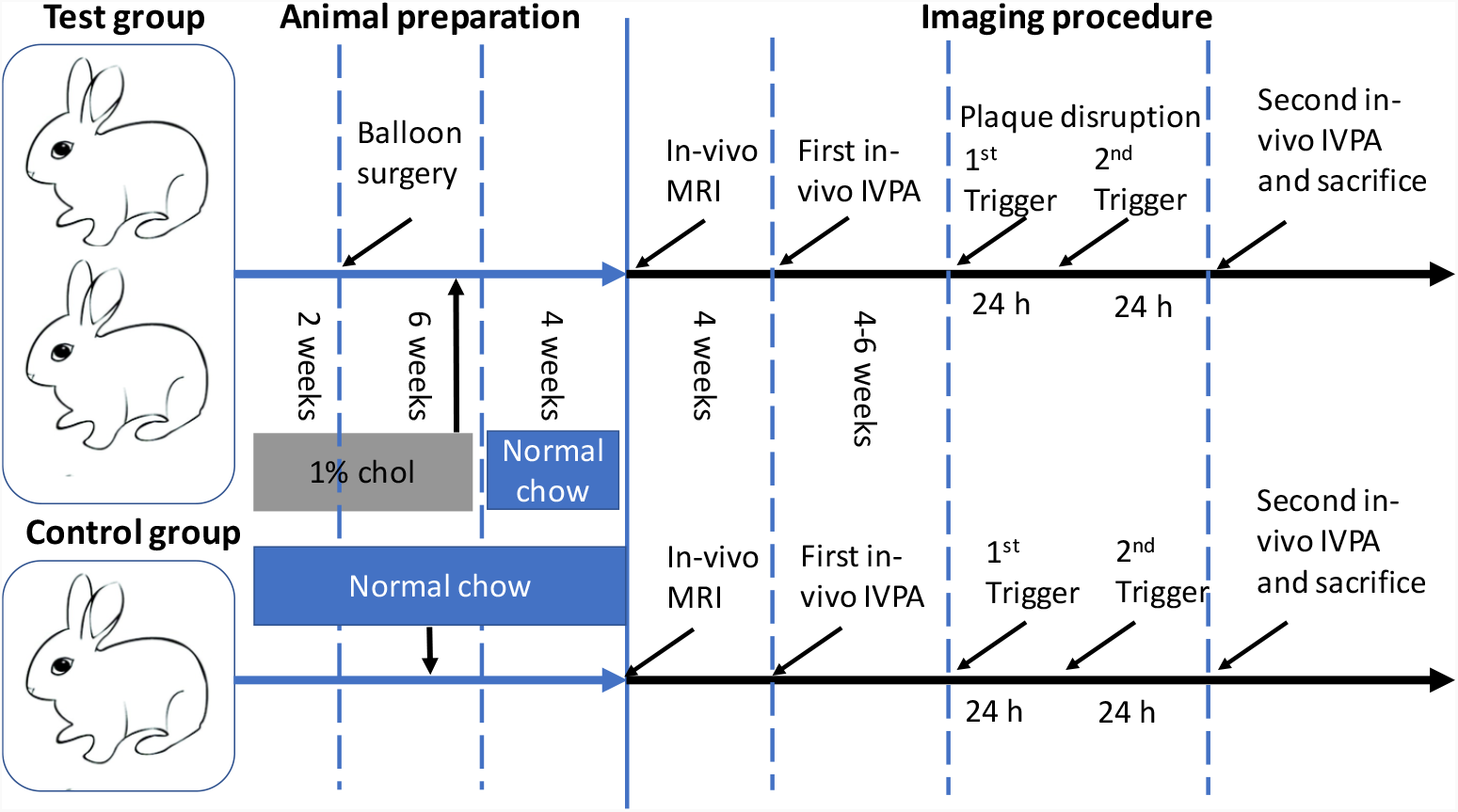
Timelines for the rabbits with or without cholesterol feeding and balloon injury. The timeline of the test group shown includes 8 weeks of 1% Chol feeding followed by 4 weeks of normal diet. Then, an endothelial injury was carried out in the test groups. For both groups, in vivo MRI was carried out at the 13^th^ week, followed by first in vivo IVPA imaging, pharmacologic triggering, and second in vivo IVPA imaging of the same plaque in the same animal. Chol: Cholesterol.

The animal protocol, under protocol #201800536, is approved by Boston University IACUC.

### The endothelial injury

The injury protocol was performed by the Hamilton lab which has extensive experience with the balloon injury model. The NZW rabbits were feed rabbit chow with 1% cholesterol for 2 weeks followed by an endothelial injury and then 6 weeks of cholesterol diet and 4 weeks of normal chow^39^. The endothelial injury of the abdominal aortic wall was performed using a 3F Fogarty catheter introduced through a right femoral artery cut down under general anesthesia. After the catheter was advanced to the diaphragm, the balloon was inflated, and the catheter was gently retracted toward the iliofemoral artery.

### In vivo MRI

In vivo MRI imaging was performed after 10 weeks of the balloon surgery. The MRI experiments on a NZW rabbit were performed under deep sedation using a 3.0-T Philips Intera Scanner (Philips Medical Systems, Ohio), offering a spatial resolution of 0.55 mm. Ungated coronal 3D phase contrast MR angiograms (PC-MRA) acquired with a T1-weighted, fast-filed echo sequence were used as scout images. Two-dimensional, T1-weighted, black-blood (T1BB) axial images were then acquired with a double-inversion recovery turbo-spin echo sequence and cardiac gating. Subsequently, ungated axial 3D PC-MRA images were acquired immediately after a bolus injection of Gd-DTPA (0.1 mmol/kg IV) (Magnevist, Germany). Finally, post-contrast-enhanced (post-CE) T1BB images were acquired 10 to 15 minutes after Gd-DTPA injection with parameters identical to those used for the non-contrast-enhanced T1BB images.

### Plaque disruption method

We attempted to disrupt the plaque by intraperitoneal injection of Russell’s viper venom (0.15 mg/kg; Enzyme Research, South Bend, IN) followed by intravenous injection of histamine (0.02 mg/kg; Sigma-Aldrich, St. Louis, MO). This procedure was performed twice within 48 h on two consecutive days. Russell’s viper venom is a procoagulant factor and endothelial toxin, and histamine acts as a vasopressor in rabbits. Heparin (1000 USP units; Sigma-Aldrich) was administered intravenously prior to euthanasia to prevent postmortem blood clotting.

### IVPA system

The IVPA system (**Figure 2a**) developed by the Cheng lab provides dual-modality intravascular photoacoustic and ultrasound imaging with real-time display^27^. A Nd:YAG pumped OPO (made by Nanjing Institute of Advanced Laser Technology) emitting ∼5 ns pulse with 500 Hz repetition rate at a wavelength of 1732 nm served as the excitation laser source. At this wavelength, the laser excites the second overtone of C-H stretch vibration in lipids. The laser output was coupled to a multimode fiber (FG200LEA, Thorlabs). The laser passes through a lab-designed fiber rotary joint and coupled to the IVPA-US catheter. At the distal end of the catheter (**Figure 2b**), the laser was reflected by a 45° gold coated rod mirror. The generated photoacoustic signal was collected by an ultrasound transducer (0.5×0.6×0.2 mm^3^, 42 MHz, 50% bandwidth, AT23730, Blatek Industries, Inc.) where ultrasound signal was converted into electronic signal. Then, signal was amplified by a pulse receiver (5073PR, Olympus Inc.) and digitalized by a data acquisition card (ATS9350 PCI express digitizer, AlazarTech). After 4 μs delay, the transducer was excited by an electronic pulse and the reflected ultrasound signal was also collected by the same ultrasound transducer, amplified and digitalized. The synchronization and delay were controlled by a function generator (9512+, Quantum Composer). All the recorded data were processed by a LabVIEW program and displayed in real time. The output pulse energy from the catheter tip was controlled around 100 μJ, corresponding to a laser fluence of 50 mJ/cm^2^ and below the ANSI laser safety standard of 1 J/cm^2^ at 1730 nm.

**Figure 2.**
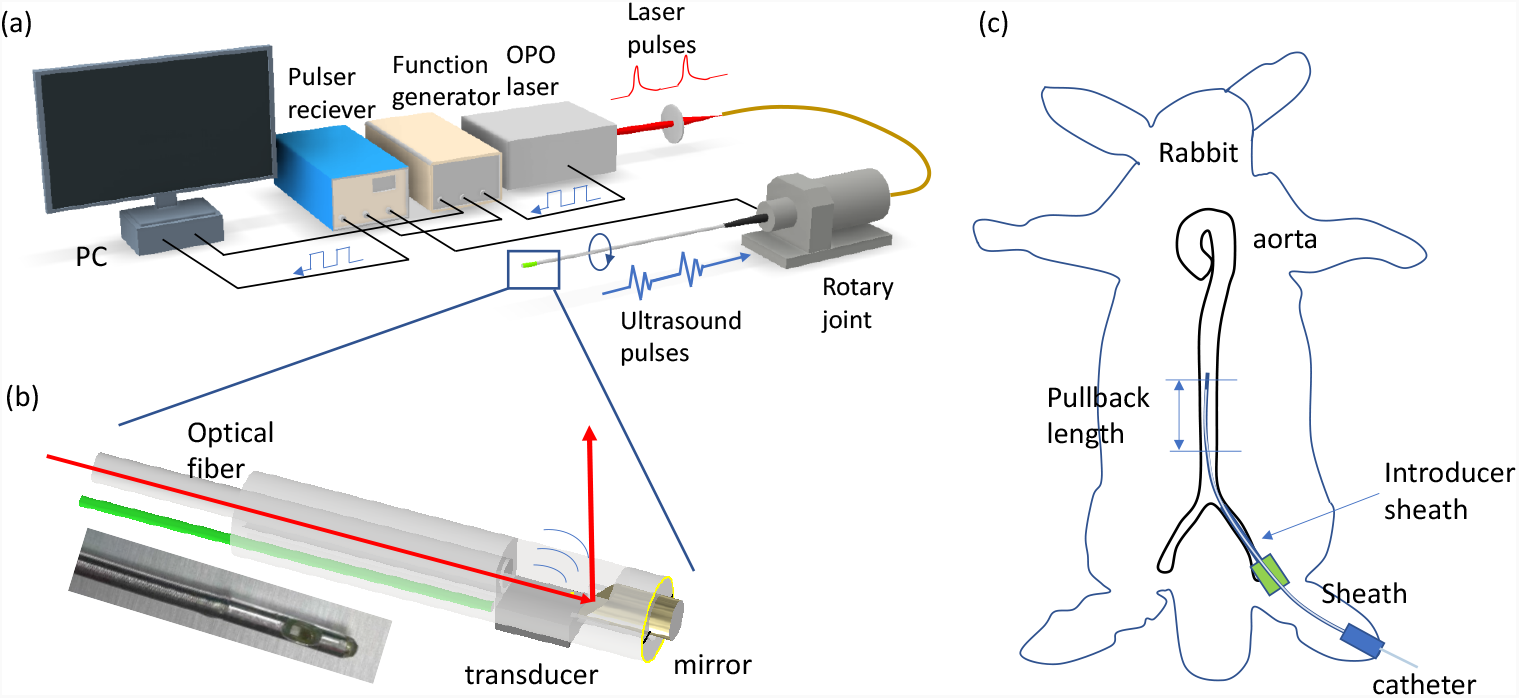
IVPA imaging system and surgical procedure for in vivo IVPA imaging. (a) The IVPA imaging system. Nanosecond laser pulses are couple to a multimode fiber and directed to the catheter through a rotary joint. At the catheter tip, the laser pulse is reflected by a gold coated mirror and shine on the artery. At the same time, the laser generates an electric trigger signal, sent to the function generator. The function generator produces two triggers, one to the data acquisition card and one to the Pulser receiver, which generates a delayed ultrasound pulse at the ultrasound transducer. (b) Schematic of catheter tip. A picture is shown below the schematic. (c) The survival surgery for in vivo IVPA imaging. An introducer catheter is inserted at the iliac artery and the catheter in sheath is inserted to the aorta.

### In vivo IVPA imaging

The survival surgery (**Figure 2c**) was conducted by Dr. Luis Guerrero from Massachusetts General Hospital. The first in vivo IVPA imaging was conducted after 10 weeks and the second one was conducted 18-20 weeks after the balloon surgery. The sedation is achieved by Ketamine/Xylazine/Acepromazine: 35 mg/kg / 5 mg/kg / 0.75 mg/kg IM. Induction of anesthesia is accomplished by injection of Acepromazine 0.75 mg/kg IM followed in at least 5 minutes by ketamine 35 mg/kg IM and Xylazine 5 mg/kg IM. A laryngeal mask (LMA) is used for the rabbit to receive oxygen from a tank.

Following induction of anesthesia, the fur on the right leg and right abdominal region connected to the leg is cleaned by a clipper. Following shaving of the site, betadine is applied, followed by alcohol, and this procedure is repeated for 3 times. A femoral artery cut down is performed under sterile conditions.

Next, a 4F introducer sheath placement in femoral artery is carried out before imaging. Through the introducer sheath, the IVPA catheter covered by catheter sheath is advanced to the thoracic aorta. The advance distance is decided by the balloon surgery area and the MRI imaging. IVPA imaging of the aorta is performed at rotation speed of 2 fps and pullback speed of 0.2 mm/s. A total length of 50 mm is recorded by pullback of the catheter. During the surgery, the animal surgeon places the looser sutures spaced further apart to lessen the number of sutures used and prevent skin irritation. IVPA imaging was performed without blood flushing. The excitation wavelength was 1730 nm and the excitation power was 30 mW at the sample.

After the first imaging procedure, the catheter and sheath are then removed. Hemostasis is accomplished by direct pressure to the artery following sheath removal and suture closure. The surgical wound is then sutured and closed. During the surgery, respiration, heart rate and SpO2 of the rabbits is monitored. PDS monofilament sutures are used to lessen the chance of post-operation irritation. Immediately after the surgery, when the animals become conscious, a plastic e-collar is placed around the neck to prevent chewing of the sutures at the incision site. All these procedures ensure the NZW survival after the imaging procedure.

After conduction of the second in vivo IVPA imaging, the animal under anesthesia is euthanized by pentobarbital overdose (>120 mg/kg intravenously). Euthanasia is accomplished by first administering acepromazine 1 mg/kg IM once. After 15-20 minutes, a small intravenous injection is placed in an ear vein, after which a lethal bolus dose of sodium pentobarbital (120 mg/kg intravenously) is administered. Cardiac arrest is confirmed with auscultation using a stethoscope.

Monitoring under anesthesia and during surgery was continuously performed by trained animal technicians. Specifically, animal respiration, heart rate, oxygen saturation, and rectal temperature were monitored by a veterinary pulse oximeter. Body temperature was maintained by a heating pad.

### Histology examination

After euthanasia, the aortas were harvested. All the aortas were fixed in 10% formalin for approximately 30 min to maintain lumen as close to in vivo morphology as possible. The aorta sections shown by IVPA imaging to have plaque were selected and cut into 3 mm segments. Then the segments were paraffin embedded, sectioned and stained with Oil Red O and H&E.

## Results

Based on the in vivo MRI measurements, we assessed the vessel wall area (VWA) and plaque gadolinium (Gd) contrast enhancement. Selected segments representing regions with or without plaque and normal vessel areas were compared. Atherosclerosis was observed in the test group rabbits and no atherosclerosis was observed in control rabbit.

Figure 3 shows representative MRI images of the aorta with plaque and the control group aorta. Contrast-enhanced MRI using gadolinium-diethylenetriamine penta-acetic acid (Gd-DTPA) has improved the discrimination between the fibrous cap and the lipid core^43,44^, and necrotic core^45^ and the visualization of coronary atherosclerosis^46^. After administration of Gd-DTPA^47^, MRI images showed hyperintense signal associated with vulnerable plaques, which are often asymmetric in the vessel wall. The presence of crescent-shaped enhancement pattern after injection of Gd-DTPA was statistically higher in vulnerable plaques. In the in vivo MRI, the test group showed a strong crescent-shaped enhancement pattern the control group had a thinner vessel wall and very low Gd-DTPA uptake.

**Figure 3.**
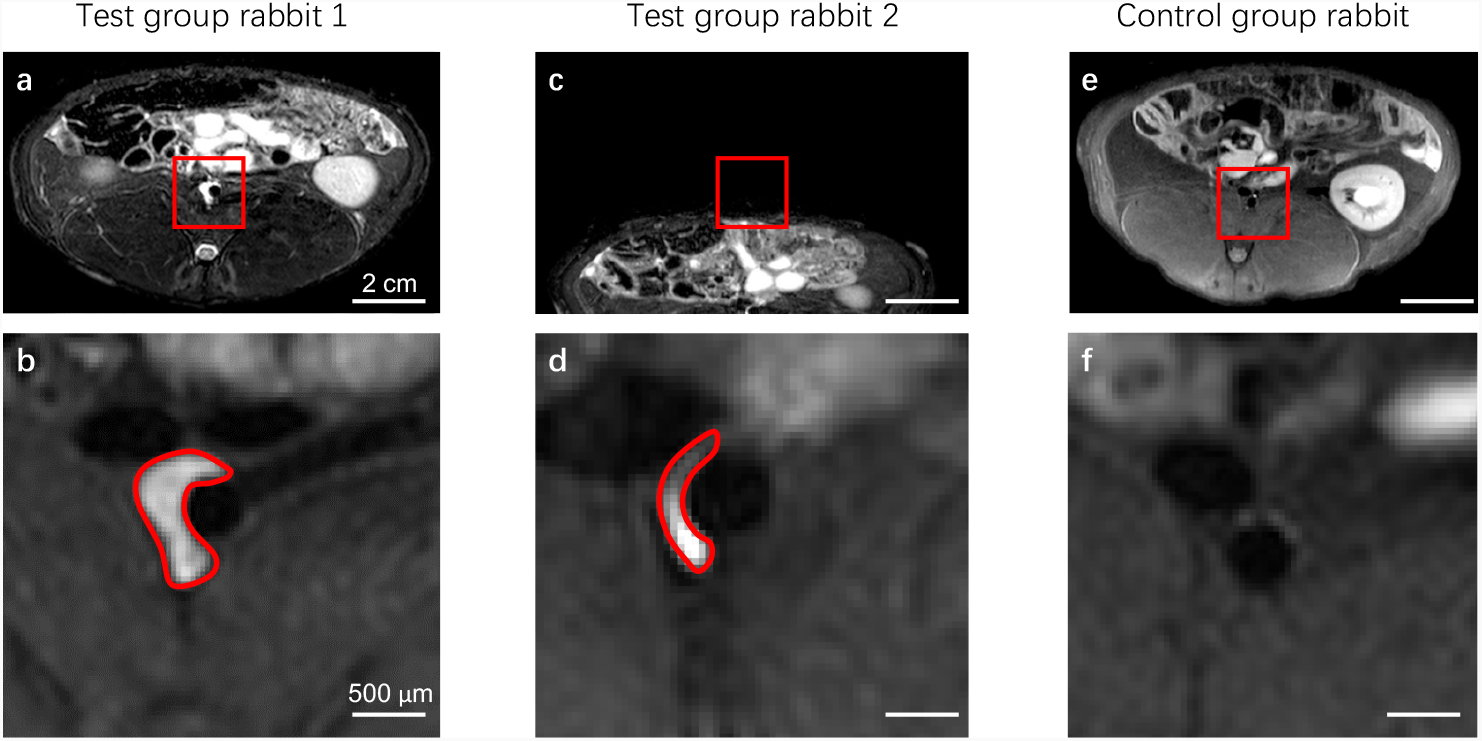
In vivo MRI images of rabbit aorta. (a,c) T1W MRI images of atherosclerosis plaque with Gd contrast enhancement in rabbit abdominal aorta taken at 10 weeks after the balloon surgery (prior to the in vivo IVPA). (e) MRI images of the control group. (b,d,f) Corresponding enlarged view of the aorta inside the red square in panels a, c and e. The areas marked by red line indicate the Gd infiltration.

Based on cross-sectional MRI along the aorta (see **videos 1, 2, 3**), we were able to determine the rough position of inflammation in the aorta. This information was used as a guide to determine the insertion length of the catheter during IVPA imaging of the plaque in the aorta.

Four weeks post MRI, we conducted survival IVPA imaging of the NZW rabbits for both test and control groups. With the laser wavelength at 1730 nm for excitation of the first overtone of C-H vibration, we recorded in vivo IVPA/US images of the aorta with 50 mm pullback length. Panels in the first row of **Figure 4** show representative cross-sectional PA images. Lipid-rich plaques are indicated by arrows. Panels in the second row of **Figure 4** show merged PA/US images. The PA images show the presence of lipid-rich plaque (red) in the lumen of the aorta wall. The US images provide morphological information about the artery. From the figure, we can see that the test group rabbits have lipid-rich plaque, and the thickness of the artery wall is much larger than the control group. In the control group, no PA signal has been observed, corresponding to absence of lipid-laden plaques. The outer vessel wall is much more detectable by US.

**Figure 4.**
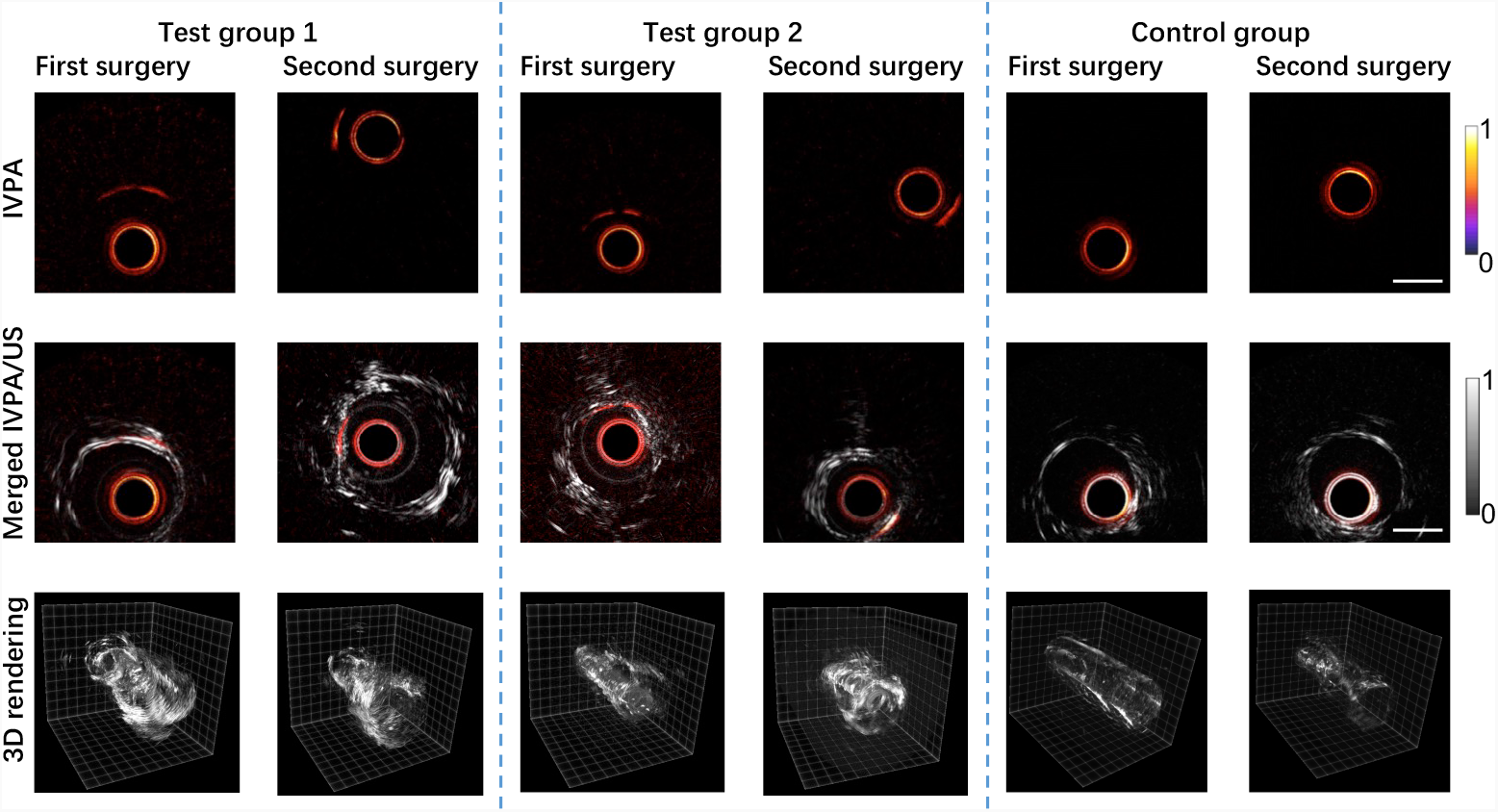
In vivo IVPA imaging of rabbit aorta. First row: Cross-sectional intravascular PA images. The red circle in the PA image originates from the photoacoustic signal generated by the sheath. Second row: Overlaid IVPA and IVUS images corresponding to the upper row. The scale bar is 1 mm for all panels. Third row: 3D rendering of the pullback IVPA images.

To quantitatively analyze the lipid core parameters before and after the pharmaceutical triggering, we generated 3D rendering of the cross-sectional IVPA images (**Figure 4**, third row). Overall, rabbits in the test group, fed on cholesterol-rich diet, showed more lipid deposition in the aorta than the rabbit in the control group.

Because IVPA imaging is able to determine the lipid amount and position at each cross section of the artery, we plotted the lipid core size and depth from the lumen with respect to the pullback positions (**Figure 5**). Over the range of 0.8 millimeters, we did not observe obvious changes in either lipid size or lipid position. These data indicate that the pharmaceutical triggering did not cause plaque rupture in the current study.

**Figure 5.**
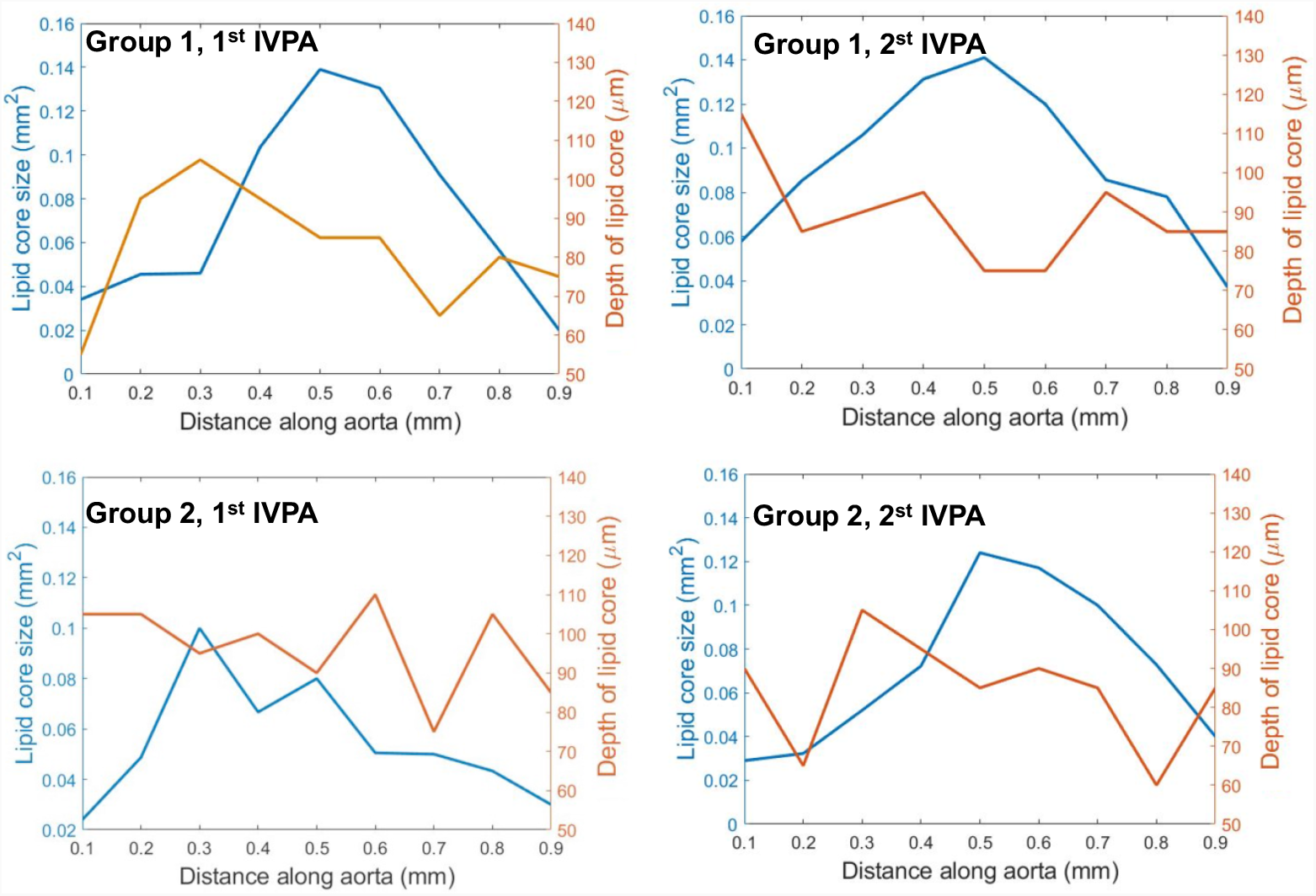
Lipid core size and depth from the lumen surface at different pullback positions for Group 1 and 2 rabbit IVPA imaging before and after the triggering.

After the second IVPA imaging, animals were euthanized and the aortas were harvested. To validate the IVPA imaging data, we stained the aorta with Oil Red O, a fat-soluble dye that stains neutral triglycerides and lipids. Meanwhile, the H&E stain provides a comprehensive picture of the microanatomy of organs and tissues. The histology shown in **Figure 6** confirmed the existence of lipid-rich plaque and the thickening of the aorta in both rabbit in the test group, but not in the control group.

**Figure 6.**
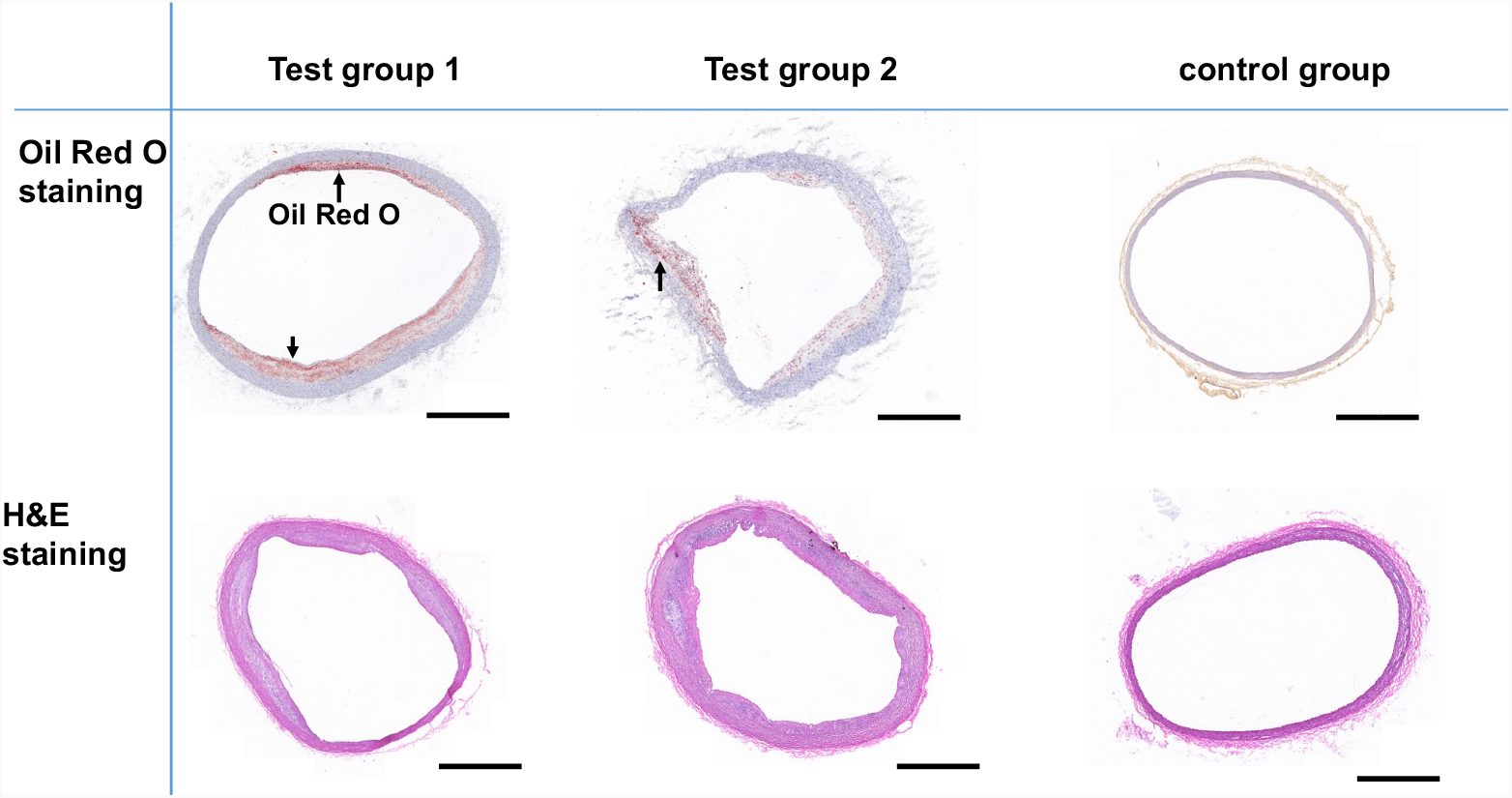
Histology of the rabbit aorta. At the end of our timeline (see Fig. 1) the rabbit was sacrificed, and the aorta cut to slices and stained with Oil Red O and H&E. The test groups showed thicker vessel walls and stronger Oil Red O staining (indicated by arrows) than the control group. Scale bar is 1 mm for all panels. Test groups 1 and 2 were balloon injured and the control group was fed normal chow and not balloon-injured.

## Discussion

There are a few challenges facing the study of thrombotic events and plaque vulnerability in live animals, including controlled induction and timing of the thrombosis. The rabbit model of controlled atherothrombosis, combining cholesterol feeding with early arterial wall injury, produces aortic plaques that mimic 4 of the 6 American Heart Association stages found in humans, including advanced highly inflammatory rupture-prone lesions in a short term (2-3 months) ^39^. The most unique and valuable contribution to atherosclerosis research is that vulnerable and stable plaques can be differentiated by a timed pharmacological injection (“triggering”) that is coordinated with vascular imaging to detect thrombus formation in a non-fatal manner ^39^. In this pilot study, we established the feasibility of monitor the same plaque in a rabbit before and after the triggering. Though we did not observe plaque rupture in the current study, the technology developed here built a foundation for future study of the relationship between vulnerability and lipid content of a plaque.

Technically our team made a few advances that enabled in vivo longitudinal IVPA imaging on the same artery in the same animal. In an earlier study, we identified a proper protective sheath material that is transparent to both PA and US signals, which is essential for in vivo applications^28^. Yet, survival IVPA imaging was not established. In this work, we further reduced the catheter diameter from 1.6 mm to 0.9 mm. Because of the smaller size of the catheter, the sheath size is reduced from 6F (1.37 mm inner diameter) to 4F (1.07 mm inner diameter). In this way, the surgical damage to the rabbit aorta is much reduced, rendering it possible to suture the incision after IVPA imaging. Furthermore, we managed IVPA imaging in a fully sterilized surgery room rather than in an optical room as in our earlier in vivo IVPA work. This procure effectively reduced the risk of infection. Finally, we applied MRI to accurately locate the plaque in the aorta. These efforts together enabled survival IVPA imaging on the same plaque before and after a treatment.

On the limitation side, the imaging speed (one cross-sectional frame per second) of our current setup is not sufficient for survival IVPA imaging of coronary artery in a swine model. This limited speed is largely due to the insufficient laser repetition rate (500 Hz). To overcome this limitation, one could use a nanosecond laser of higher pulse energy and higher repetition rate (2 kHz), which has allowed fast IVPA imaging at the speed 16 frames per second ^27^.

In summary, using a sheathed IVPA catheter and a rabbit aorta balloon injury model, aided with in vivo MRI, we demonstrated the feasibility of survival IVPA/US imaging of the same lipid-laden plaque in the same animal. The reported imaging system and method promise a valuable asset for understanding the progression of the atherosclerosis and evaluating the effectiveness of a medical treatment.

## Acknowledgments

The authors acknowledge Dr. Luis Guerrero for performing the survival surgery and thank Yingchun Cao for help and discussion. We also thank Raisa Khuda for assistance in preparation of the manuscript.

## Author contributions

Yi Zhang and Erik Taylor carried out the IVPA imaging experiments. Nasi Huang carried out the MRI experiment. James Hamilton and Ji-Xin Cheng guided the project. Yi Zhang and Ji-Xin Cheng co-wrote the manuscript. All authors read the manuscripts.

**Additional Information Competing financial interests: J.-X.C. has a financial interest in Vibronix Inc, which does not support this work**.

